# Femtomolar and selective dopamine detection by a gold nanoparticle enhanced surface plasmon resonance aptasensor

**DOI:** 10.1101/273078

**Authors:** Yong Cao, Mark T. McDermott

## Abstract

Ultrasensitive and selective detection and quantification of dopamine (DA) plays a key role in monitoring neurodegenerative diseases. However, the detection limit reported for DA detection is typically in the lower nM range. Pushing the detection limit to pM or lower for this particular target to cover the physiological levels (< 130 pM) is significant. Herein, DA DNA aptamer (DAAPT) gold nanoparticle (AuNP) conjugate is utilized to enhance the surface plasmon resonance (SPR) signal, which enables to detect and quantify DA in the femtomolar (200 fM) to picomolar range. To the best of our knowledge, this is the lowest detection limit achieved for SPR sensing of dopamine. The as-prepared 10 nm DAAPT-AuNP conjugate demonstrates strong binding affinity (K_d_ = 3.1 ± 1.4 nM) to the complementary DNA (cDNA) probe on gold chip. The cDNA probe is immobilized to the chip via polydopamine surface chemistry, which allows the Michael addition of any primary amine-terminated biomolecules. By adjusting the concentration of the DAAPT-AuNP conjugate, two calibration curves are generated with dynamic ranges from 100 µM to 2 mM, and from 200 fM to 20 nM, respectively. Both calibration curves have negative slopes, showing good agreement to a dose-response curve in an enzyme inhibition assay. In addition, the sensing strategy is evaluated to be specific for DA detection using a series of DA analogs and other metabolites as potential interferences.

## 1. Introduction

Small-molecule metabolites (MW < 1000 Da), the intermediates and products of metabolism of living organisms, are now serving as an important type of biomarker, thanks to the rapid development of metabolomics [1,2]. Metabolites sensing plays a crucial role in using the information for medical diagnostics. Dopamine is a neurotransmitter that is a member of the catecholamine family in the brain and a precursor to epinephrine and norepinephrine [3]. Also, it is a major transmitter in the extrapyramidal system of the brain, important in regulating movement, and related to neurodegenerative diseases, such as Parkinson’s disease, Alzheimer’s disease, and Huntington’s disease [4,5]. Sensing strategies that can provide fast, ultrasensitive and selective analysis of dopamine is key in monitoring these neurodegenerative diseases. Sensitive and selective analysis of small-molecule metabolite in general is of great importance in human health monitoring.

The physiological levels of dopamine vary in different human biofluids (blood, urine, cerebrospinal). According to the Human Metabolome Database (HMDB, http://www.hmdb.ca/), the concentration of dopamine (HMDB0000073) in blood is less than 130 pM, while in human cerebrospinal fluid and urine the levels of dopamine are ∼5 nM and less than 1 μmole/mmole creatinine. The reported limit of detection value for dopamine is typically in the nM range or higher (μM), either based on electro-chemical-based [6-12] or optical-based [13-15] methods. Compared the reported limit of detection values to the above physiological levels (especially in blood), there still exists strong motivation to push the detection limit to lower pM range or even fM. Therefore, methods/assays that can detect and quantitate dopamine to cover the whole physiological levels are highly demanded.

Surface plasmon resonance (SPR) is an optical sensing platform that allows for label-free, sensitive and quantitative detection of biomolecules [16]. A variety of SPR-based assays/sensors have been reported to measure proteins [17-19] and nucleic acids [20]. Detecting small molecules using this technique is a little bit trickier than proteins and nucleic acids, due to the smaller size and lower molecular weight of this type of targets. However, researchers have incorporated a variety of nanomaterials [21, 22] such as metal nanoparticles [23-25], magnetic nanoparticles [26,27], quantum dots [28], and carbon-based nanomaterials [29,30] into the assays to enhance the signal and lower the detection limits. For example, Corn et al reported the employment of nucleic acid functionalized nanoparticle-enhanced SPR imaging to measure protein and DNA targets down to the 25 fM based on oligonucleotide and aptamer microarrays [31]. We reported a small-molecule functionalized gold nanoparticle enhanced competitive assay to detect and quantitate folic acid in the nM range with a detection limit of 2.9 nM [32]. Zhou et al reported the use of gold nanoparticles-aptamer conjugates as a competitive reagent to detect adenosine down to 0.1 nM [33]. Split aptamer together with gold nanoparticle enhancement was utilized to detect adenosine with a limit of detection of 1.5 pM [34]. Based on these work, we believe a nanoparticle-enhanced SPR assay combined with highly specific recognition element could push the detection limit for dopamine to lower pM or fM, thus close the gap mentioned above.

In this work, we incorporate dopamine DNA aptamer (DAAPT) [35,36] conjugated gold nano-particle (AuNP) to detect and quantitate dopamine down to 200 fM using SPR. We first prepared the 10 nm DAAPT-AuNP conjugate, which is quite stable in buffer solution. Since the DAAPT can bind to dopamine with high affinity, the whole conjugate is “OFF” in the presence of DA, and is “ON” in the absence of DA. On the chip surface, a complementary single-stranded DNA (cDNA) probe is immobilized that can hybridize to DAAPT. The polydopamine surface chemistry enables the attachment of the amine-terminated cDNA probe via Michael addition reaction. When the DAAPT-AuNP probe is “OFF” (DA present), it cannot bind to the cDNA probe on chip surface, thus no signal will be observed. On the contrary, the DAAPT-AuNP probe that is “ON” (DA absent) will bind to the cDNA probe, generating a big signal intensity. Based on this strategy, a negative correlation is created between the SPR signal response and the concentration of DA. The as-prepared 10 nm DAAPT-AuNP conjugate probe shows strong binding affinity (K_d_ = 3.1 ± 1.4 nM) to the cDNA probe. By changing the concentration of the DAAPT-AuNP conjugate, we performed the inhibition assay to detect and quantitate DA in two different concentration ranges: 100 µM – 2 mM and 200 fM – 20 nM. Both calibration curves have a negative slope, show good sensitivity and reproducibility. Moreover, our assay is evaluated to be specific for DA analysis, with little interference observed from DA analogs and other metabolites.

## 2. Experimental

### 2.1 Chemicals and reagents

10 nm (5.7 × 10^12^ particles/mL, 9.47 nM), 20 nm (7.0 × 10^11^ particles/mL, 1.16 nM), 30 nm (2.0 × 10^11^ particles/mL, 0.33 nM), and 40 nm (9.0 × 10^10^ particles/mL, 0.15 nM) of citrate-capped gold nanoparticle stock solutions were purchased from Ted Pella, Inc. Deionized (DI) water (H_2_O) with a resistivity greater than 18 MΩ was filtered in a Barnstead Nanopure purification system. 10 × PBS (phosphate buffered saline, 1× = 10 mM phosphate, 0.154 M NaCl) buffer, dopamine (DA) hydrochloride, folic acid (FA), 17β-estradiol (E2), L-3,4-dihydroxyphenylalanine (DOPA), 3,4-dihydroxyphenylacetic acid (DOPAC), 1,2-dihydroxybenzene (Catechol), epinephrine (EP), homovanillic acid (HVA), L-ascorbic acid (AA) were all purchased from Sigma-Aldrich and used as received. Tris-HCl buffer (10 mM, pH 8.5) was obtained from bioWORLD of GeneLinx International, Inc. 20× TBS (tris buffered saline, 1× = 25 mM Tris, 0.15 M NaCl) was purchased from ThermoFisher Scientific. SF-10 glass sheet (100 mm × 100 mm × 1 mm) was purchased from Schott Glass. (Tridecafluoro-1,1,2,2-tetrahydrooctyl)-1-dimethylchlorosilane was purchased from United Chemical Technologies. All DNA oligos at 250 nmole with HPLC purification were ordered from Integrated DNA Technologies, Inc. The sequence information of all single-straded DNA oligos are as follows: a thiol-terminated dopamine DNA aptamer (DAAPT) (5’-S-S(CH_2_)_6_-GTC TCT GTG TGC GCC AGA GAC ACT GGG GCA GAT ATG GGC CAG CAC AGA ATG AGG CCC-3’), an amine-terminated complementary DNA (cDNA) probe (P20) (5’-NH_2_(CH_2_)_12_-GGG CCT CAT TCT GTG CTG GC-3’), and an amine-terminated negative control DNA probe (A20) (5’-NH_2_(CH_2_)_12_-AAA AAA AAA AAA AAA AAA AA-3’).

### 2.2 Fabrication of SPRi chip with multiple Au spots

The purchased SF-10 glass sheet (100 mm × 100 mm × 1 mm) was cut into small square substrate (18 mm × 18 mm × 1 mm). The substrates were cleaned using hot piranha solution (1:4 30% H_2_O_2_:H_2_SO_4_) followed by thorough rinsing with DI water. Substrates were blown dry using Ar gas. [*Warning: Piranha solution should be handled with extreme care; it is a strong oxidant and reacts violently with many organic materials. It also presents an explosion danger. All work should be performed under a fume hood with appropriate personal safety equipment*.] The cleaned and dried substrates were mounted to a mask that expose nine 2 mm diameter round spots and placed into the chamber of a thermal evaporator (Torr International Inc., New Windsor, NY). Metal films of chromium (2 nm) and gold (42 nm) were sequentially coated on the glass substrate through the mask. After removal from the evaporator, the substrates were then exposed to a vapor of (tridecafluoro-1,1,2,2-tetrahydrooctyl)-1-dimethylchlorosilane under reduced pressure for 24 h to create a hydrophobic background on the glass surface. The SPRi chips were stored in a desiccator under vacuum at room temperature until use.

### 2.3 Dopamine DNA aptamer-gold nanoparticle (DAAPT-AuNP) conjugate preparation

10 µL of 100 µM of the thiol-terminated DAAPT solution (PBS) was added to 1mL of 10 nm, 20 nm, 30 nm, 40 nm citrate-capped gold nanoparticle stock solutions, respectively. The mixed solutions were incubated at room temperature for 24 h. Next, 500 µL of PBS buffer (pH 7.4, 0.154 M NaCl) was added to each mixed solution, which was incubated at room temperature for another 24 h. The 10 nm, 20 nm, 30 nm, and 40 nm mixed solutions were centrifuged (Eppendorf a5417R microcentrifuge) under 14000 rpm for 45 min, 13000 rpm for 30 min, 10000 rpm for 30 min, and 10000 rpm for 30 min, respectively. The supernatant of each solution was carefully removed to avoid any loss of the particles. The AuNP pellet was re-dispersed in 1 mL DI H_2_O, or PBS buffer, or TBS buffer, and stored at 4 °C in the dark until use.

### 2.4 UV-vis spectroscopy

Extinction spectra of 10 nm, 20 nm, 30 nm, and 40 nm citrate-capped AuNP stock solutions and corresponding DAAPT-AuNP conjugate solutions in different media (DI H_2_O, PBS, TBS) were obtained by UV-vis spectroscopy. 500 µL of each AuNP solution was added to a quartz micro-cuvette for UV-vis measurement. All solutions were measured in transmission mode in a double-beam Perkin-Elmer Lambda 35 instrument. For each measurement, the corresponding medium used for re-dispersing gold nanoparticle pellet was used as the reference.

### 2.5 SPRi measurement of 10 nm DAAPT-AuNP conjugate binding to cDNA probe

Prior to the modification, the homemade SPRi chip was rinsed with pure ethanol and DI water and dried under an argon gas flow. Polydopamine (PDA) surface chemistry described elsewhere was used to immobilize cDNA probe (P20) to the gold chip surface [37]. Briefly, a 2.5 µL droplet of 10.5 mM DA in Tris buffer (pH 8.5) was added to each gold spot of the chip. The chip with the solution droplets was stored in a humid petri dish for 10 min at room temperature. Then, the chip was rinsed with DI water and dried under argon gas flow. Each spot of the chip was next exposed to a 2.5 µL droplet of 250 µM of amine-terminated cDNA (P20) probe or negative control DNA (A20) for 12 h in the petri dish. The surface was then blocked by 1 mg/mL ethanolamine solution for 1 h. After rinsing and drying, the modified chip was mounted to the SPRi instrument (Horizon SPRimager, GWC Technologies, Madison, MI) to measure the binding of 10 nm DAAPT-AuNP conjugate to the cDNA probe. The apparatus has been described in detail elsewhere.[38, 39] 10 nm DAAPT-AuNP conjugate in PBS buffer solutions at 6 concentrations (0.095 nM, 0.24 nM, 0.47 nM, 0.95 nM, 2.4 nM, 4.7 nM) were prepared from the stock DAAPT-AuNP solution (9.5 nM). Each solution was exposed to the modified chip surface, and the real-time SPRi sensorgram was recorded and analyzed.

### 2.6 Inhibition Assays

All assays were carried out on chips containing cDNA (P20) and negative control (A20) spots. As described below, two concentration ranges of DA were studied. For the higher (µM to mM) range, 500 µL of 9.5 nM 10 nm DAAPT-AuNP conjugate solution (in PBS buffer) was mixed with 500 µL of DA standard solution (in PBS buffer) at a specific concentration (100 µM, 200 µM, 300 µM, 500 µM, 1 mM, 2 mM) for 1 h. Each mixed solution was flowed into the instrument and exposed to gold chip surface modified with P20 and A20 DNA probes, and then a real-time SPRi sensorgram was obtained for each mixed solution. For the lower (fM to nM) range, we decreased the concentration of the 10 nm DAAPT-AuNP conjugate solution to 1.9 nM. The NP solution was mixed with DA standard solutions (200 fM, 300 fM, 500 fM, 2 pM, 20 pM, 20 nM, 20 µM) for 1 h. Each solution was flowed into the instrument and the SPRi sensorgram was obtained.

### 2.7 Specificity studies

To test the specificity of the assay, we chose a variety of DA analogs (DOPA, DOPAC, Catechol, EP, HVA, AA) and other two metabolites (FA, E2). 500 µL of 1.9 nM 10 nm DAAPT-AuNP conjugate solution (in PBS buffer) was mixed with 500 µL of 20 nM of the possible interference standard solution described above for 1 h. Each mixed solution was measured using SPRi, and the result was compared with that for DA.

## 3. Results and discussion

Ultrasensitive and selective detection of DA as well as other critical neurotransmitters plays a significant role in monitoring some neurodegenerative diseases, such as Parkinson’s, Alzheimer’s, and Huntington’s diseases, as mentioned above. Herein, we have developed an aptamer-gold nanoparticle conjugate-enhanced inhibition assay using surface plasmon resonance imaging that enables ultrasensitive and selective detection of DA down to the fM concentration range. First, we attach the bio-recognition element – a DNA aptamer for DA (DAAPT) – to gold nanoparticle (AuNP) surface, and obtain this critical DAAPT-AuNP conjugate. The sensing strategy of the DAAPT-AuNP conjugate-enhanced inhibition assay is shown in Fig. 1. In the absence of DA, no binding happened between DAAPT on AuNP surface and DA, the conjugate probe is in its “ON” state. With a partially complementary single-stranded DNA (cDNA) probe on the SPRi chip surface, we can detect the binding of the conjugate to the cDNA probe. As a result, a big signal response will be generated. On the contrary, in the presence of a higher concentration of DA, all the DAAPTs on the AuNP conjugate surface will bind to their target, so the conjugate probe is in its “OFF” state. In this case, the conjugate will not bind to cDNA on chip surface due to the blocking of the DNA aptamer by the binding of DA, thus no signal response will be observed. During the whole process, the target DA molecule serves as a “key” to turn “ON” and “OFF” the DAAPT-AuNP conjugate. In theory, if we fix the initial concentration of the conjugate, the effective concentration of the probe that can bind to cDNA surface will decrease with increasing DA concentration. Therefore, the signal response is inversely related to the concentration of DA, and we will see a negative slope for the calibration curve.

**Figure 1.**
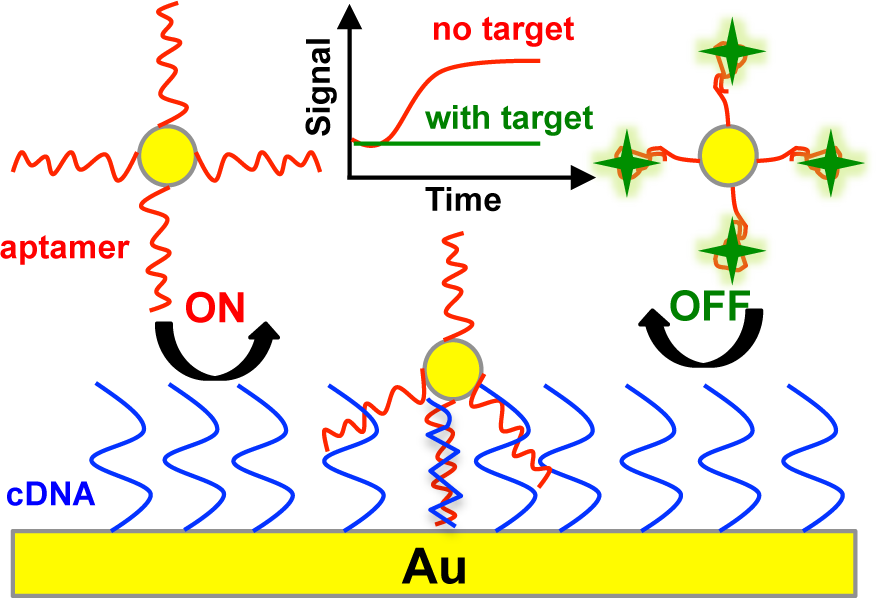
Schematic illustration of the DNA aptamer-gold nanoparticle conjugate enhanced sensing strategy for dopamine (DA) analysis using SPR imaging. Amine-terminated single-stranded complementary DNA (cDNA) to DNA aptamer is immobilized on gold chip surface via polydopamine surface chemistry. In the absence and presence of DA, the conjugate is in its ‘ON’ and ‘OFF’ states respectively. ‘ON’ state means the conjugate can bind to the immobilized cDNA, thus generate a big SPR signal. On the contrary, ‘OFF’ state means no binding will occur between the conjugate and the cDNA due to the blocking of DNA aptamer by the binding of DA.

We prepared 10 nm, 20 nm, 30 nm, and 40 nm DAAPT-AuNP conjugate probes via the strong Au-S interaction [40,41] between the thiol-terminated DAAPT and AuNP. Fig. 2A shows the extinction spectra of 10 nm citrate-capped AuNP in H_2_O and 10 nm DAAPT-AuNP conjugate in three different media (H_2_O, PBS, TBS). All the spectra have a good shape with the localized surface plasmon resonance (LSPR) peaks located around 520 nm. Taking a closer look at the amplified portion around the maximum extinction of the spectra (Fig. 1B), we can observe clear red shifts of the LSPR peaks from 517 nm (citrate-capped, H_2_O) to 525 nm (DAAPT-AuNP, H_2_O), to 524 nm (DAAPT-AuNP, PBS/TBS). The observed red shifts of the LSPR peaks confirm the successful conjugation of DAAPT to AuNP surface. Moreover, the good shape of the spectra in PBS/TBS means the prepared 10 nm DAAPT-AuNP conjugate is quite stable in buffer solution with high salts concentration. The stability of 10 nm DAAPT-AuNP conjugate in buffer solution is key for the success of the inhibition assay, because the conjugate will mix with DA standard solution or unknown sample solution with a relatively high salt environment. Extinction spectra for 20 nm, 30 nm, and 40 nm DAAPT-AuNP conjugates are shown in Fig. S1. Compared to 10 nm DAAPT-AuNP conjugate, 20 nm and 30 nm conjugates do show red shifts of the LSPR peaks, but they are not stable in buffer solution. For 40 nm conjugate, no red shift of the LSPR peaks is observed. Based on these studies, we chose 10 nm DAAPT-AuNP conjugate and used it in the competitive assay. It has been reported that spherical nucleic acid gold nanoparticle conjugate can be formed by densely functionalizing the gold cores with a surface shell of DNA coordinated via sulfur groups to the gold, and the size of the gold core in the original work is 13 nm [42,43].

**Figure 2.**
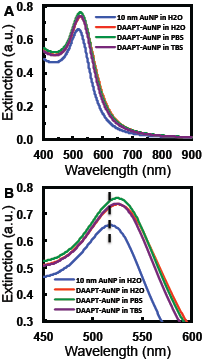
UV-vis characterization of the as-prepared 10 nm dopamine aptamer-gold nanoparticle (DAAPT-AuNP) conjugate in H_2_O and buffers. (A) Extinction spectra for as-purchased 10 nm citrate-AuNP in H_2_O (blue) and as-prepared 10 nm DAAPT-AuNP conjugate in H_2_O (red), PBS (green), and TBS (purple). (B) Amplified portion around the maximum extinction for clarity. The dashed line marks the maximum extinction of 10 nm citrate-AuNP.

**Figure 3.**
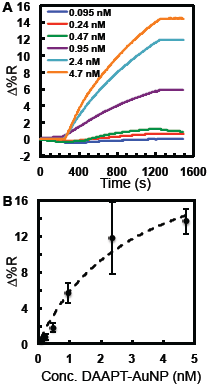
Binding of 10 nm DAAPT-AuNP conjugate to cDNA probe on gold chip surface. (A) Typical SPRi sensorgrams for the binding of 10 nm DAAPT-AuNP at six concentrations: 0.095 nM (blue), 0.24 nM (red), 0.47 nM (green), 0.95 nM (purple), 2.5 nM (cyan), and 4.7 nM (orange). (B) A plot of SPR signal vs. DAAPT-AuNP solution concentration for adsorption to cDNA. The symbols are the data points and the dashed line is the least-squares fit of Langmuir adsorption isotherm equation (R^2^ = 0.9669). The error bars represent standard deviation for triplicate measurements.

To immobilize the cDNA probe to the SPRi chip surface, we chose polydopamine (PDA) surface chemistry that allows the Michael addition of amine-terminated cDNA molecule to PDA film on chip surface [44,45]. The protocol reported elsewhere [37] was followed, which have approved the successful attachment of amine-terminated DNA molecule to the gold spot surface. The SPRi sensorgram for the growth of PDA film on gold spots surface is shown in Fig. S2. The binding of 10 nm DAAPT-AuNP conjugate to cDNA probe on chip surface was investigated by SPRi measurement. Fig. 2A contains the SPRi sensorgrams for 10 nm DAAPT-AuNP conjugate binding under different concentrations: 0.095 nM, 0.24 nM, 0.47 nM, 0.95 nM, 2.4 nM, and 4.7 nM. The SPRi signal intensity increases with increasing 10 nm DAAPT-AuNP conjugate concentrations. The least concentrated (0.095 nM) and most concentrated (4.7 nM) 10 nm DAAPT-AuNP conjugate solutions give 0.4 ± 0.1 and 13.6 ± 0.4 of change in percent reflectivity (Δ%R), respectively. By comparing the two signals, we conclude that the signal response is coming from specific binding of 10 nm DAAPT-AuNP conjugate to the cDNA probe. Isotherm was constructed for 10 nm DAAPT-AuNP conjugate and cDNA binding from sensorgrams shown in Fig. 2A. The 10 nm conjugate binding data points were fitted with a Langmuir isotherm (Fig. 2B) equation using the one-site ligand-binding model included in SigmaPlot (Systat Software, Inc., San Jose, CA). The binding data agree well (R^2^ = 0.9669) with the Langmuir isotherm, and the dissociation constant (K_d_) and the maximum signal intensity (Δ%R_max_) for the binding were determined to be 3.1 ± 1.4 nM and 23.8 ± 5.6, respectively. It was reported in the literature that the binding constant between spherical nucleic acid functionalized gold nanoparticle conjugate and its complementary oligonucleotide sequence in a homogeneous solution can be down to the picomolar range, or even lower, depending on the density of the spherical nucleic acid on nanoparticle surface and the length of the complementary nucleic acid [46,47]. The binding constant value reported here is a few orders of magnitude higher than the literature values, probably because our assay is heterogeneous and the complementary DNA probe is immobilized on a solid surface. This may introduce more steric effect and hindrance to the binding, which results in a lower binding affinity.

To detect and quantitate DA, the 10 nm DAAPT-AuNP conjugate solution at a fixed concentration was mixed with DA standard solution at a specific concentration. By pre-mixing the two solutions, DA molecules will bind to DAAPT-AuNP conjugates, thus decreasing the effective concentration of DAAPT-AuNP conjugate that can bind to cDNA probe on chip surface. Here, we can play with two crucial factors: the concentration of 10 nm DAAPT-AuNP conjugate as well as the concentration of DA standard solution. As stated previously, the signal intensity should decrease with increasing DA concentration in the mixed solution. Also, the dynamic range of the calibration curve can be easily adjusted by simply changing the concentration of 10 nm DAAPT-AuNP conjugate solution. We started with the most concentrated (9.5 nM, before mixing) 10 nm DAAPT-AuNP conjugate solution, and mixed it with DA standard solutions ranging from 100 µM to 2 mM in a 1:1 volume ratio. Some representative SPRi sensorgrams for the exposure of these mixed solutions to cDNA on chip surface are shown in Fig. 4A. As expected, we see a decrease of signal response with increasing DA concentration in the mixed solutions. The signal intensities for the least concentrated (100 µM) and the most concentrated (2 mM) DA solutions are 13.91 ± 2.69 and 1.15 ± 0.14, respectively. The signal intensity for the most concentrated is approximately 8% of that for the least concentrated. The calibration curve (Fig. 4B), a plot of signal response vs. dopamine concentration, demonstrates this trend with a negative slope. Also, the calibration curve exhibits good sensitivity and reproducibility, which enables detecting of DA ranging from 100 µM to 2 mM. To further decrease the dynamic range, we lowered the concentration of 10 nm DAAPT-AuNP conjugate to 1.9 nM. Fig. 5A shows some representative SPRi sensorgrams for the exposure of mixed solutions of 1.9 nM (before 1:1 mixing) 10 nm DAAPT-AuNP conjugate solution and DA standard solutions ranging from 0.2 pM to 2 µM. Similarly, the signal decreases from 5.64 ± 0.20 (blank) to 1.70 ± 0.33 (2 µM). The calibration curve (Fig. 5B) has a negative slope, good sensitivity, and reproducibility, showing a two-orders-of-magnitude dynamic range from 0.2 pM to 20 pM. Last but not least, the specificity of the proposed assay for DA detection was studied by choosing a series of DA analogs and other metabolites, including DOPA, DOPAC, catechol, epinephrine (EP), homovanillic acid (HVA), ascorbic acid (AA), folic acid (FA), and 17β-estradiol (E2). Some of the analogs get involved in the metabolic pathway for dopamine metabolism as well, thus are good candidates for assay specificity studies [3,48,49]. For each possible interference, DA standard solution in the mixed solution containing 1.9 nM (before mixing) DAAPT-AuNP conjugate was replaced to one of these DA analogs or metabolites at a certain concentration, and exposed to cDNA on SPRi chip surface. The results are summarized in Fig. 6. A big signal decrease was observed for mixed solution containing DA. Minor signal decrease is observed for DOPA, Catechol, EP, and FA. However, the magnitudes of signal decrease for DOPA and EP are almost negligible when compared to that for DA. This gives us confidence that the proposed assay is specific for DA analysis, with little interference from its analogs and other metabolites.

**Figure 4.**
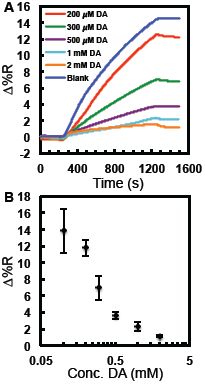
Detection of DA using the inhibition strategy from higher micromolar (µM) range to lower milimolar (mM) range. (A) Representative SPRi sensorgrams for mixed solutions (1:1 volume ratio) of 10 nm DAAPT-AuNP solution at fixed concentration (9.5 nM, before mixing) and DA standard solutions at different concentrations: blank (no DA, blue), 200 µM (red), 300 µM (green), 500 µM (purple), 1 mM (cyan), and 2 mM (orange). (B) Calibration curve (SPR signal vs. DA concentration) for the determination of DA in the corresponding concentration range. The error bars represent standard deviation for triplicate measurements.

**Figure 5.**
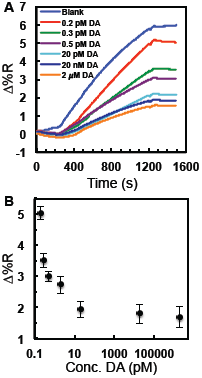
Detection of DA using the competitive assay from higher femtomolar (fM) range to lower picomolar (pM) range. (A) Representative SPRi sensorgrams for mixed solutions (1:1 volume ratio) of 10 nm DAAPT-AuNP solution at fixed concentration (1.9 nM, before mixing) and DA standard solutions at different concentrations: blank (no DA, blue), 200 fM (red), 300 fM (green), 500 fM (purple), 20 pM (cyan), 20 nM (dark blue), and 2 µM (orange). (B) Calibration curve (SPR signal vs. DA concentration) for the determination of DA in the corresponding concentration range. The error bars represent standard deviation for triplicate measurements.

**Figure 6.**
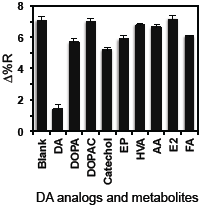
Specificity study of the competitive assay using DA analogs (DOPA, DOPAC, catechol, epinephrine, homovanillic acid, ascorbic acid) and other metabolites (estradiol and folic acid) as potential interferences. The error bars represent standard deviation for at least triplicate measurements of each small molecule.

## Conclusions

In summary, dopamine DNA aptamer (DAAPT) conjugated gold nanoparticle (AuNP) is utilized for ultrasensitive and selective detection and quantitation of DA down to 200 fM using surface plasmon resonance (SPR) imaging. This is the lowest detection limit achieved for SPR sensing of dopamine. By pre-incubating the 10 nm DAAPT-AuNP conjugate with DA, the conjugate probe can be turned “ON” and “OFF” by controlling the concentration of DA. A negative correlation between SPRi signal intensity and DA concentration is established based on this inhibition detection format. The as-prepared 10 nm DAAPT-AuNP conjugate probe shows high binding affinity to the partially complementary DNA probe on chip, with a dissociation constant (K_d_) equal to 3.1 ± 1.4 nM. Two calibration curves were generated with dynamic ranges from 100 µM to 2 mM, and from 200 fM to 20 pM, respectively, which suggests that the dynamic range can be adjusted by changing the concentration of 10 nm DAAPT-AuNP conjugate probe. In addition, the proposed assay exhibits good sensitivity, reproducibility, and high specificity for DA detection. Furthermore, this nanoparticle enhanced SPR aptasensor can potentially be applied to other analytes (either proteins or small molecules) as well, as long as the target has a corresponding aptamer.

## Acknowledgements

The authors would like to thank the NanoFab at University of Alberta for facility use. This work was supported by Alberta Innovate Health Solutions (AIHS) Collaborative Research and Innovation Opportunities (CRIO) grant and the Natural Sciences and Engineering Research Council (NSERC) of Canada (Discovery Grant to MTM).

## Appendix A. Supplementary data

Supplementary data associated with this article can be found, in the online version, at Extinction spectra for 20 nm, 30 nm, and 40 nm citrate-AuNPs and DAAPT-AuNPs in different media; SPRi sensorgram for the growth of polydopamine film on the gold chip surface.

## Table of Contents (TOC) Figure

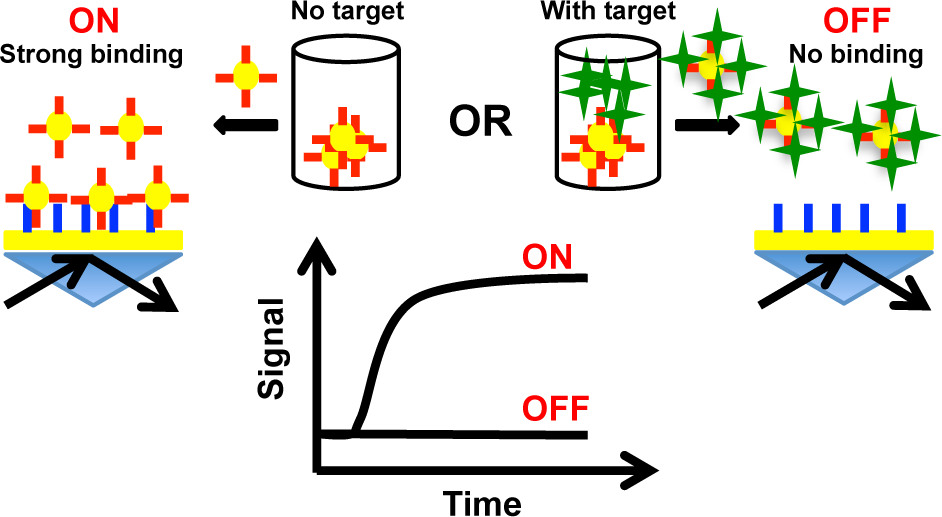

